# The bat influenza H17N10 can be neutralized by broadly-neutralizing monoclonal antibodies and its neuraminidase can facilitate viral egress

**DOI:** 10.1101/499947

**Authors:** George Carnell, Efstathios S Giotis, Keith Grehan, Francesca Ferrara, Stuart Mather, Eleonora Molesti, Simon Scott, Antonello Pessi, Krzysztof Lacek, Nigel Temperton

## Abstract

The diversity of subtypes within the Influenza A virus genus has recently expanded with the identification of H17N10 and H18N11 from bats. In order to further study the tropism and zoonotic potential of these viruses, we have successfully produced lentiviral pseudotypes bearing both haemagglutinin H17 and neuraminidase N10. These pseudotypes were shown to be efficiently neutralized by the broadly-neutralizing monoclonal antibodies CR9114 and FI6. Our studies also confirm previous reports that H17 does not use sialic acid as its cellular receptor, as pseudotypes bearing the H17 envelope glycoprotein are released into the cell supernatant in the absence of NA. However, we demonstrate that N10 facilitates heterosubtypic (H5 and H7) influenza HA-bearing pseudotype release in the absence of another source of NA, significantly increasing luciferase pseudotype production titres. Despite this, N10 shows no activity in the enzyme-linked lectin assay used for traditional sialidases. These findings suggest that this protein plays an important role in viral egress, but is perhaps involved in further accessory roles in the bat influenza lifecycle that are yet to be discovered. Thus we show the lentiviral pseudotype system is a useful research tool, and amenable for investigation of bat influenza tropism, restriction and sero-epidemiology, without the constraints or safety issues with producing a replication-competent virus, to which the human population is naïve.

**Significance statement:** Influenza virus is responsible for mortality and morbidity across the globe; human populations are constantly at risk of newly emerging strains from the aquatic bird reservoir which harbors most of the subtypes of influenza A (H1-H16). Recently identified subtypes (H17N10, H18N11) from bats have broadened the reservoir from which potential pandemic strains of influenza can emerge. To evaluate the potential for these novel subtypes to cross over into human populations, their ability to establish an infection, in addition to the extent of cross-reactive immunity established by human seasonal strains needs to be investigated. This study highlights a novel platform for the study of the bat H17 and N10 envelope glycoproteins, using a lentiviral pseudotype system. Following the generation of this pseudotype it was employed in cell entry and microneutralization assays. These showed that two well-characterised monoclonal antibodies (mAb) which target avian and human influenza subtypes will also neutralize H17. Furthermore the data presented in this study show a novel aspect of the N10 glycoprotein in its ability to facilitate the budding of pseudotypes bearing different influenza HAs.

## Introduction

Influenza A virus is the principal causative agent of influenza, which is a substantial burden to global economies and represents a significant public health risk worldwide (Ma et al. 2009; Reperant et al. 2012). The first stages of the viral life cycle are mediated by the influenza glycoprotein hemagglutinin (HA), which is located in the lipid outer membrane of the virus. This protein mediates both the attachment of the influenza virus to the target cell and the fusion process which allows the virus to infect its host cell. Through its activity, HA is a key determinant of virus tropism (Dos Reis et al. 2011; Sahini et al. 2010). Furthermore, HA is a highly variable protein and its features are often used as methods for distinguishing different viral strains (Ebrahimi et al. 2014; Wilson and Cox 1990).

While the association between influenza viruses and wild birds has long been established (Gelfond et al. 2009; Webster et al. 1992), the discovery of novel influenza-like viruses in New World bats (H17N10, H18N11) represents a possible challenge to the notion that avian species are its sole reservoir (Freidl et al. 2015; Sun et al. 2013; Tong et al. 2012, 2013). Recently, Egyptian *Rousettus aegyptiacus* bats have been reported to harbor a third bat influenza subtype (Kandeil et al. 2018). It is highly likely that further subtypes will be discovered in future years, warranting further research on existing strains of bat influenza.

The H17N10 and H18N11 subtypes of influenza A were discovered in bat species in Peru and Guatemala respectively. They are significantly diverged from other known influenza strains, in respect to all eight gene segments (Tong et al. 2012, 2013). The HA of these novel strains contain a number of apparently unique structural features and exhibit receptor-binding activities that differ from other influenza viruses (Sun et al. 2013). Notably, it has been shown that H17 and H18 hemagglutinins do not bind to sialic acid receptors, and the true receptors are currently still unknown (Tong et al. 2013).

The neuraminidase (NA) of these viruses is also divergent from other known NAs, but the overall structure is preserved, despite differences in the active site (Juozapaitis et al. 2014; Ma et al. 2015). Lack of sialidase activity has been reported previously for these NAs (García-Sastre 2012; Li et al. 2012; Tong et al. 2013; Zhu et al. 2012), suggesting they utilize a different substrate altogether. The combination of atypical HA binding profiles alongside a lack of NA activity suggests that this virus functions differently to previously discovered influenza A subtypes, despite relative phylogenetic relatedness.

Although the receptor remains unknown, H17 and H18 sequences have been extensively analyzed and compared to those of other HA gene sequences, to investigate the potential for these bat viruses becoming zoonotic (Freidl et al. 2015; Mänz et al. 2013).

At present, the degree to which the H17N10 and H18N11 isolates are capable of infecting non-bat hosts is unknown, as attempts to isolate wildtype virus have not been successful. Sequence analysis indicates that there is significant potential for a spillover occurrence, but further research is required to assess the true potential of these viruses as pandemic threats (Freidl et al. 2015; Juozapaitis et al. 2014).

To study the zoonotic potential of H17N10 and H18N11 viruses, attempts to integrate bat HAs into influenza A reverse genetics systems were made but none yielded infectious virus (Juozapaitis et al. 2014; Zhou et al. 2014). More recently, progress has been achieved by Moreira et al. 2016 to this end, using a pseudotyped VSV platform.

Influenza pseudotyped viruses (PVs) are useful tools to study both viral entry mechanisms and the antibody response directed against the influenza HA and NA. Furthermore, when use of PVs is coupled with detailed sequence analysis and phylogenetic inference, they offer the potential to establish safe and effective assays to inform epidemiological and public health models of viral spread and risk. This is especially the case with later generation, single cycle lentiviral vectors, allowing experimentation on functional glycoproteins with the use of a reporter incorporated in the lentivirus genome.

In this study, we report generation of the first H17 and H17N10 pseudotyped lentiviruses, their use in virus neutralization assays using broadly neutralizing monoclonal antibodies (bnmAbs) and investigation into the debated substrate specificity of the putative N10 neuraminidase.

## Results

### Generation of H17- and N10-pseudotype viruses

Generation of H17- (A/little yellow shouldered bat/060/2010) and H5-(A/Vietnam/1194/2004) lentiviral PVs was achieved after transfection of HEK293T/17 cells with lentiviral pseudotype production plasmids (Figure 1), followed by transduction of target human U87 MG (glioblastoma) cells with PV supernatant. Generation of H17-PVs was only possible in the presence of HAT or TMPRSS2 proteases in HEK293T/17 cells (Figure 2), as has been seen with other subtypes of influenza (Bertram et al. 2010; Böttcher et al. 2006). Subsequent experiments were carried out using canine MDCK II and RIE1495 target cells which were previously reported to allow production of VSV-H17 (Moreira et al. 2016). In initial experiments, attempts were made to generate functional H17 and H17N10 PV utilizing different protease-expressing plasmids to promote HA maturation. PV luciferase-based titres increased significantly when HAT-, TMPRSS2- and TMPRSS4-expression plasmids were used in the transfection mix when compared to controls, indicating activation of the bat HA and fusion competence. Other proteases tested (furin, KLK5, and TMPRSS3) did not yield significant titre increases. MDCK II and RIE1495 cells were notably transduced, whereas HEK293T/17 cells were not. In the absence of co-transfected, protease-encoding plasmids, no significant PV titre was measurable (Figure 3).

**Figure 1.**
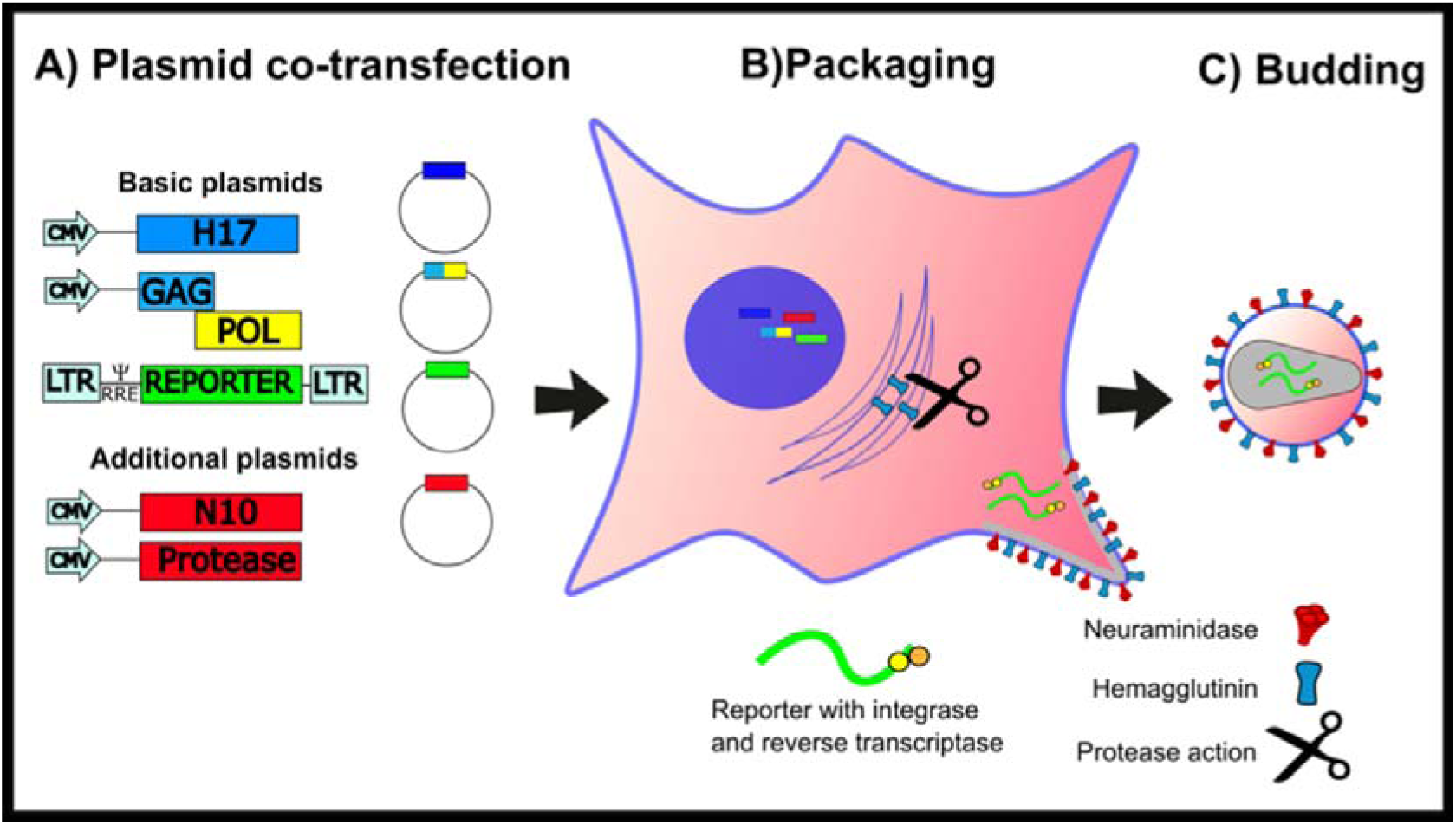
Cartoon showing the production of H17N10 PV via plasmid transfection.

**Figure 2.**
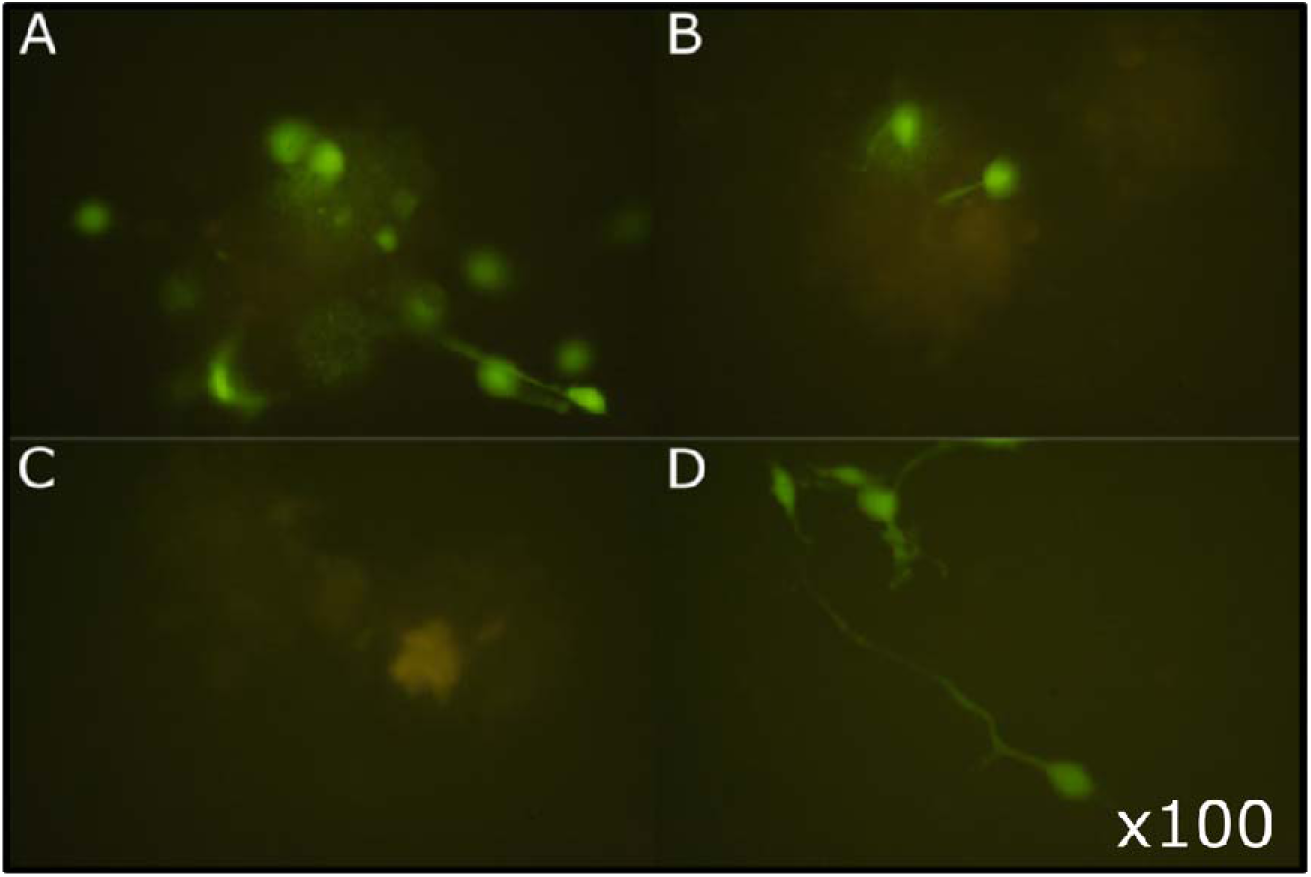
Transduction of U87 MG cells by influenza GFP PV (x100 magnification). A) H17 produced using pCAGGS-HAT. B) H17 PV produced using pCAGGS-TMPRSS2. C) Cell only control. D) H5 (A/Viet nam/1194/2004) PV positive control.

**Figure 3.**
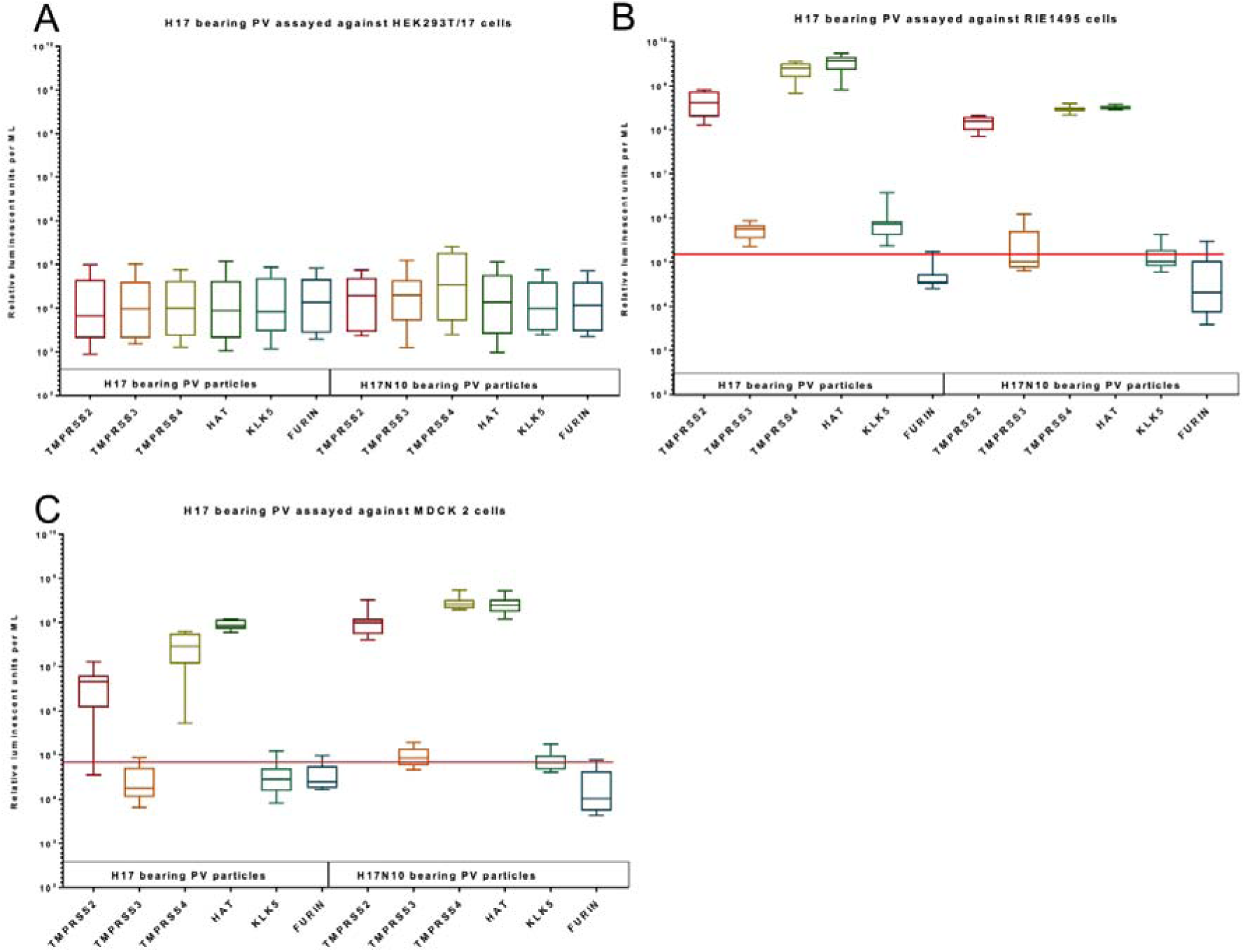
Transduction of various target cell lines. A) HEK293T/17, B) RIE1495 and C) MDCK II with H17 and H17N10 pseudotyped viruses carrying the luciferase reporter gene. Results given in Relative Luminescence Units per ml; RLU/ml). Average cell only luminescence shown as a red line.

### H17 entry of target cells is inhibited by broadly neutralizing monoclonal antibodies

Treatment of RIE1495 cells with the acidifying agent ammonium chloride resulted in significantly lower levels of luciferase activity, demonstrating that H17 requires a low pH for membrane fusion similar to conventional influenza viruses, such as H5 (Figure 4). Several studies have demonstrated that the HA trimers of conventional influenza viruses must simultaneously coordinate their conformational changes to complete membrane fusion (Otterstrom et al. 2014). HA-binding broadly neutralising antibodies (bnmAbs) typically recognize loop regions surrounding the receptor binding site or conserved regions of the stem and their induced-inhibition is serotype-specific (Dreyfus et al. 2013; Ekiert et al. 2009; Sui et al. 2009). In order to test whether cross reactive antibody responses would affect the H17 glycoprotein, several characterised bnmAbs were employed. Neutralization potency was measured as IC_50_ (concentration or serum dilution required to neutralise 50% of input virus). Neutralization tests were conducted using both the permissible target cell lines identified above (MDCK II and RIE1495). The first bnmAb, CR9114, binds to a conserved epitope in the HA stalk of group 1 and 2 influenza A viruses, and has also been shown to protect against lethal influenza challenge in a mouse model against both lineages of influenza B (Dreyfus et al. 2012). The second, CR6261, has been shown to neutralize H1 and H5 subtypes (Friesen et al. 2010). The third, FI6-nt, can neutralize H1 to H16 subtypes of influenza A. A fourth, FI6-chol is a derivative of F16-nt, conjugated to cholesterol (Corti et al. 2011; Lacek et al. 2014). H17 is effectively neutralized by CR9114, at a concentration of 0.05 μg/ml. FI6-nt and F I6-chol also neutralize H17N10-PV, however, concentrations required exceed those of CR9114. In contrast, CR6261 was not able to neutralize H17 using the H17 PV assay (data not shown). Neutralization data is shown in Figure 5.

**Figure 4.**
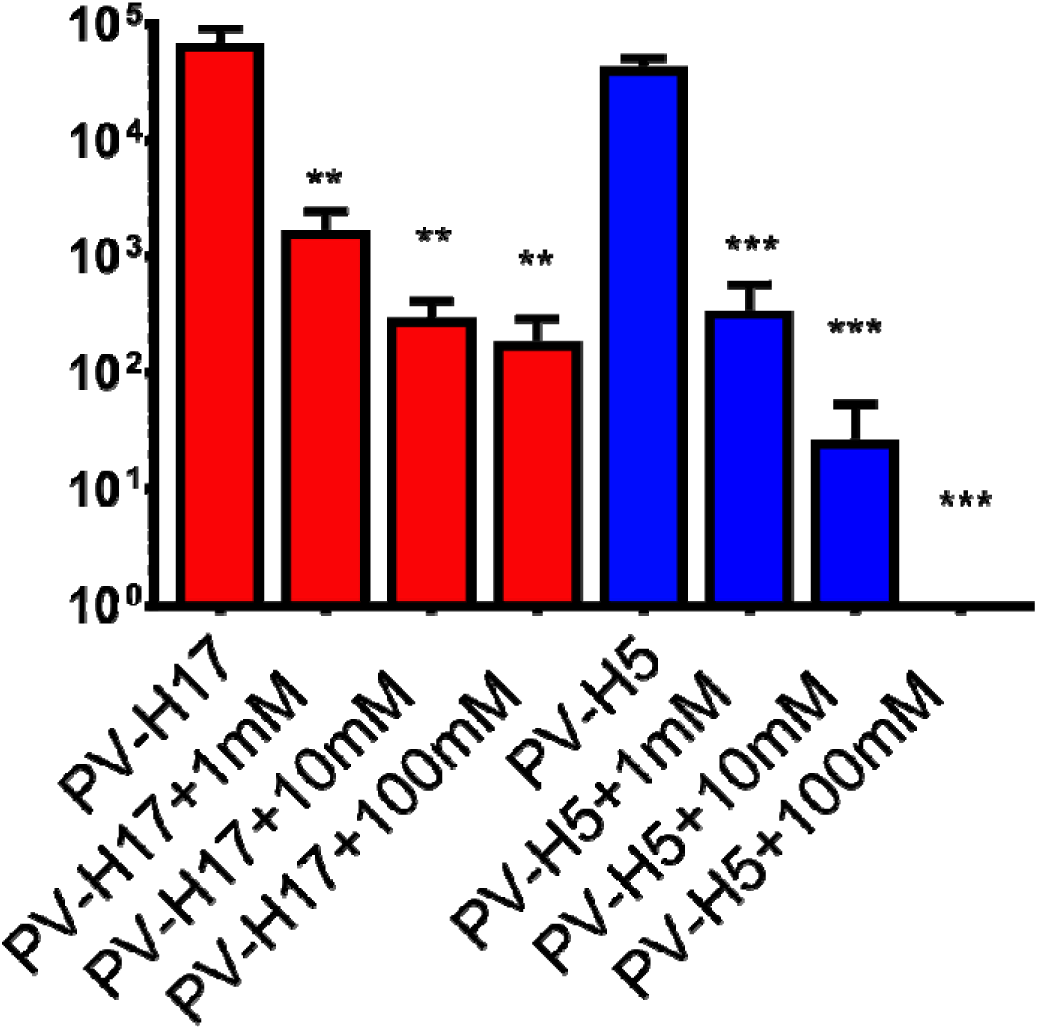
The effect of ammonium chloride based acidification on the transduction of RIE1495 cells by H17 PV. Increases in acidification of RIE 1495 cell cultures prior to transduction by H17 resulted in lower luciferase based titres, indicating that the pH sensitivity of H17 is similar to that of H5. Significant differences in transduction denoted by asteriks, ** represents p= <0.001 and *** p= <0.0001 respectively.

**Figure 5.**
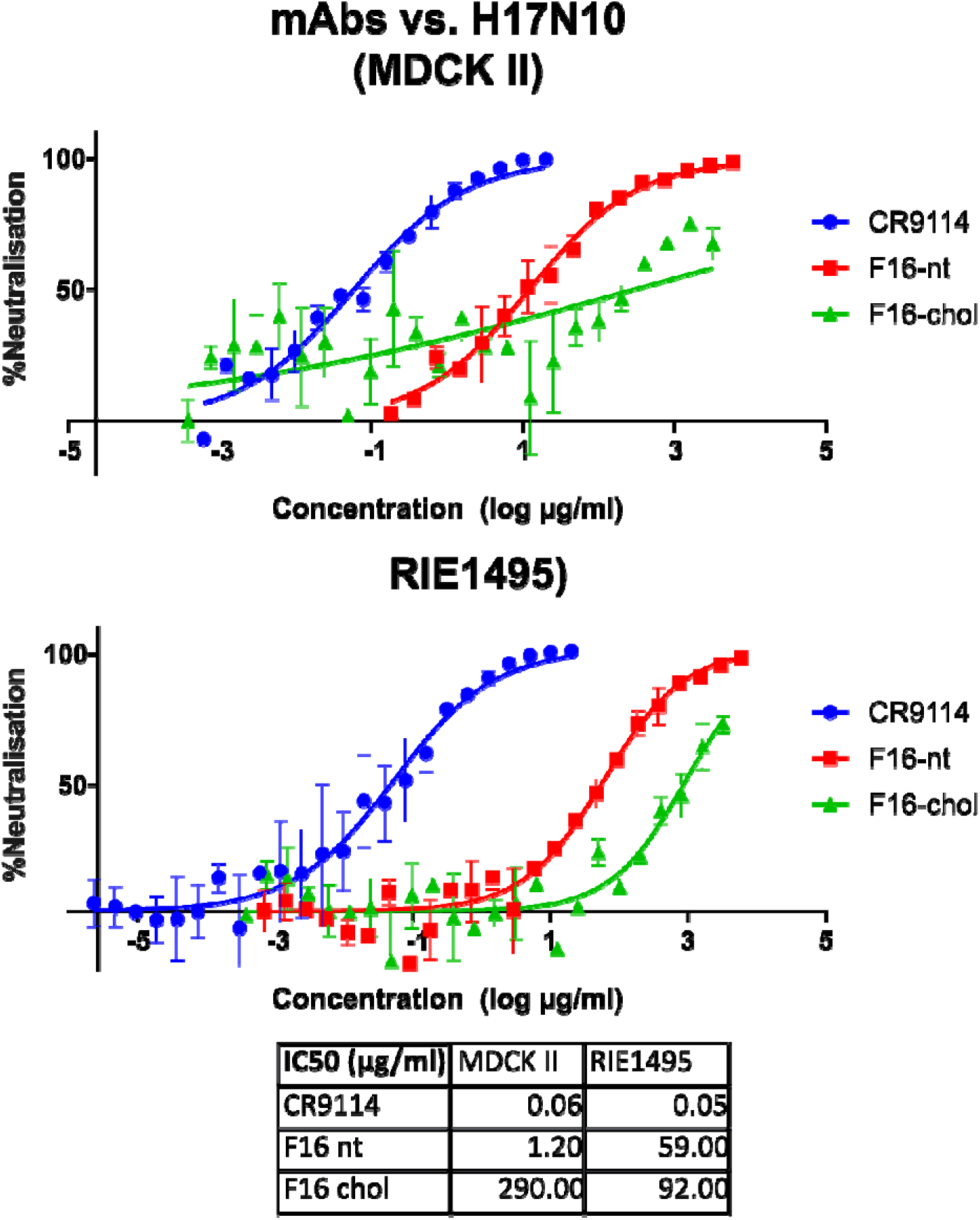
Neutralization curves and IC50 values for mAbs CR9114, FI6-nt and FI6-Chol against FI17N10 bearing lentiviral pseudotypes on cell lines RIE1495 and MDCK II. Nonlinear regression carried out using Graphpad (Prism 7) in order to provide IC_50_ values for each graph. Each IC_50_ is the concentration of mAh required for reduction of 50% of the virus input in terms of luciferase activity.

### N10 facilitates production of conventional H5 and H7 bearing PV

Pretreatment of target cells with sialidase, which removes cell surface sialic acids, did not affect the entry of H17 and H17N10 PVs, which supports previous studies that H17 does not bind sialic acids for infection (Maruyama et al. 2016), see Figure 6. N10-bearing PVs were titrated alongside H17 and H17N10 PV, in order to measure their ability to transduce cells. Results showed that N10 had no effect on H17 mediated viral entry and did not improve transduction when co-expressed with H17 (data not shown). However, addition of N10 in the generation of H5 or H7 PVs (H5N10, H7N10) produced high titre PV in the absence of any other neuraminidase source (Figure 7), indicating that N10 is facilitating release of viral particles bearing sialic acid-binding glycoproteins. H5N10 luciferase titres were one log lower than parental H5 PV produced with exogenous neuraminidase, but two logs higher than N10, ΔEnvelope (ΔEnv) or cell only controls (Figure 7). Despite this, the same H5N10 PV preparation did not show any detectable NA activity when titrated using the enzyme linked lectin assay (ELLA), confirming previous reports that this protein is not a sialidase (García-Sastre 2012; Li et al. 2012; Tong et al. 2013; Zhu et al. 2012). Similarly, multiple different preparations of H17N10 or of N10 bearing PV produced negative results in ELLA (data not shown).

**Figure 6.**
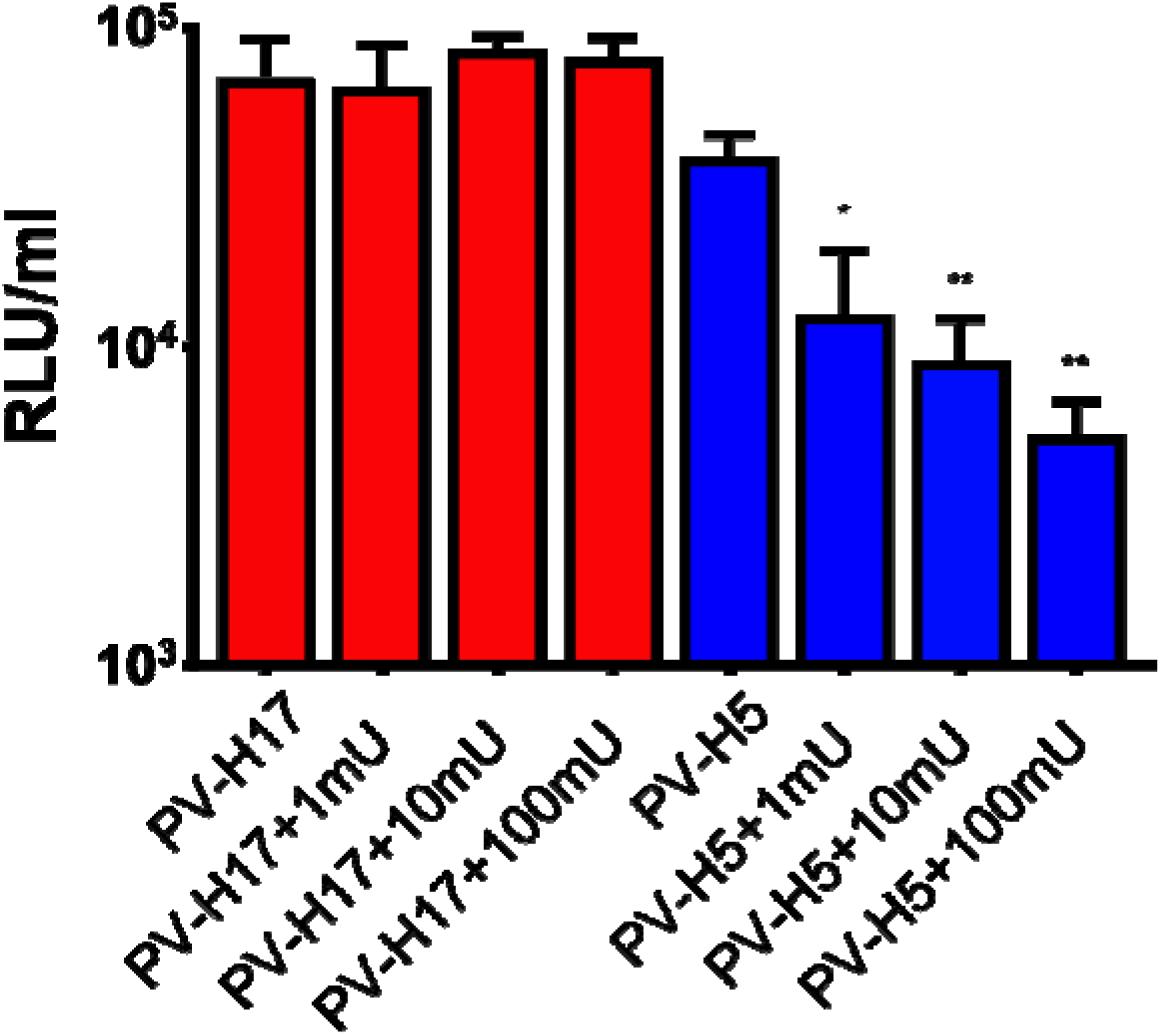
The effect of neuraminidase pre-treatment on the transduction of RIE1495 cells by H17 and H5 PV. No relationship is seen between pre-treatment of cells with neuraminidase for H17 bearing PV, but H5-based transduction is significantly reduced when sialic acids are stripped from target cells. Significant difference in transduction denoted by asteriks, * represents p= <0.01 and ** p= <0.001 respectively.

**Figure 7.**
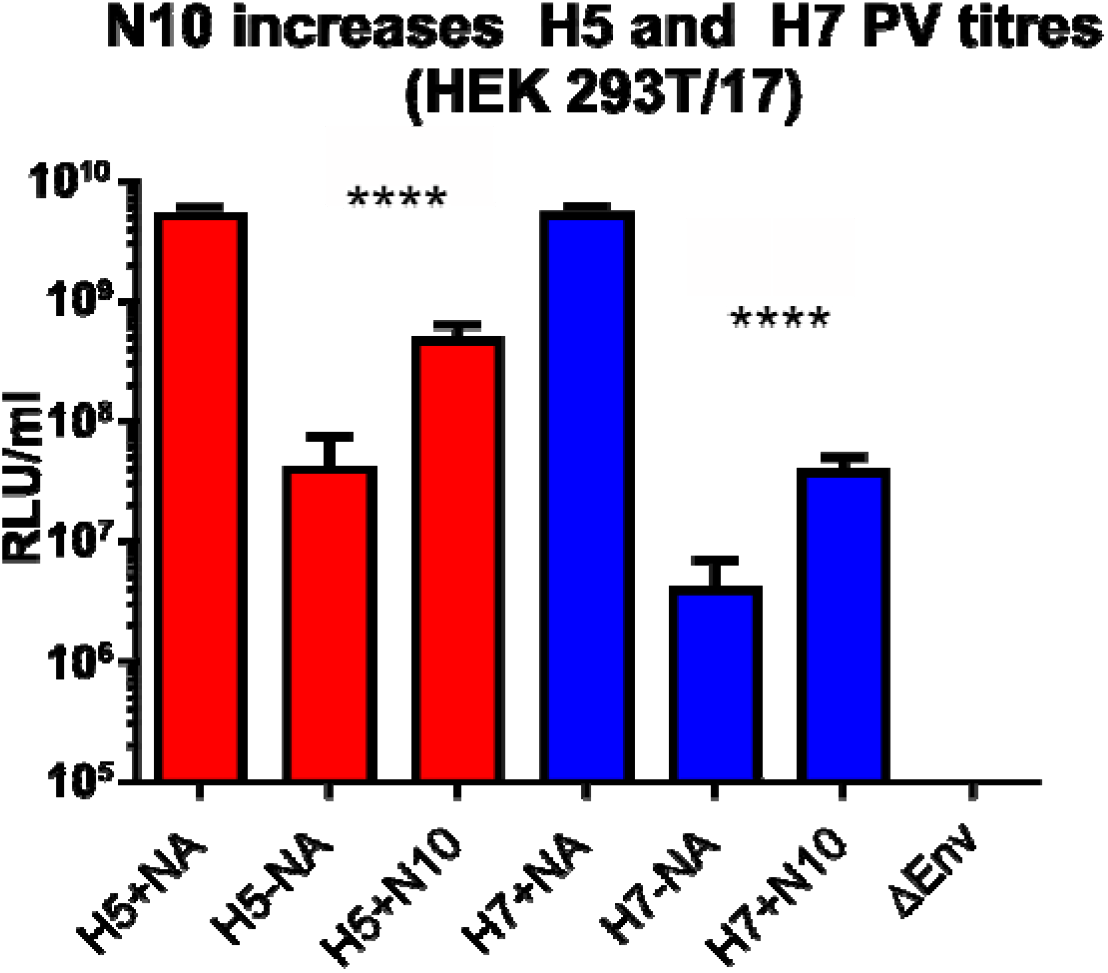
Release of H5 and H7 bearing PV by N10. PV bearing the glycoproteins H5 (A/Vietnam/1194/2004) and H7 (A/Shanghai/1/2013) were produced with exogenous neuraminidase (+NA), with no NA (-NA) or accompanied with an N10 glycoprotein (+N10). H5 and H7 PVs produced in the presence of N10 have significantly higher luficerase titres than those produced with no NA. Background luciferase titre control (Δ Env) control shown. Significant differences in transduction denoted by asteriks, **** represents p= <0.00001.

## Discussion

We have successfully produced H17 and H17N10 bearing lentiviral pseudotypes, and shown that N10 is not required for H17 pseudotype budding, but that it can mediate release of a heterologous, sialic acid-binding HA bearing PVs. These bat influenza PVs were neutralized by cross-reactive bnmAbs, suggesting that the stalk region of the H17 glycoprotein retains conserved epitopes present in group 1 and 2 influenza HAs (Sun et al. 2013).

Of the target cell lines tested, H17 and H17N10 lentiviral PV were able to transduce U87 MG, MDCK II and RIE1495 cells to varying degrees when activated by proteolytic cleavage. Reports differ on the permissibility of the MDCK I type cells (Hoffmann et al. 2016; Moreira et al. 2016). Due to the nature of the pseudotype based microneutralisation (pMN) assay used in this study, the addition of cells in suspension into PV containing supernatant may allow infection/transduction to occur before cell adhesion and thus by a different route than *in vivo.* This may explain our results in the context of Moriera and colleagues’ findings that the bat influenza H18 VSV pseudotypes enter at the basolateral membrane and inefficiently infected confluent cell monolayers (Moreira et al. 2016). MDCK I and II differ in passage number (low and high, respectively), differences between them include the mucin-type transmembrane protein podoplanin, which is expressed only in MDCK I, and the Forssman glycosphingolipid, which is expressed only in MDCK II cells (Hansson et al. 1986; Zimmer et al. 1997). The susceptibility of MDCK cells to bat influenza viruses is unclear, and is partly compounded by the availability of various lineages with different characteristics (Dukes et al. 2011). The widespread use of MDCK cells in influenza research is at odds with a previous study that showed H17 failed to bind to MDCK cells (Sun et al. 2013). While the reasons for this are currently unclear, it has been hypothesized that the level of HA binding is below the threshold of detection for the assays used in this earlier study (Maruyama et al. 2016). Alternatively, only a subset of MDCK lineages may be susceptible to these viruses (Table 2). RIE1495 cells are morphologically similar to MDCK II cells and express the Forssman antigen, a possible factor in their susceptibility to H17 bearing PVs (Moreira et al. 2016). The cell tropism data reported in the Moreira study raise some interesting ideas concerning the H17 receptor, as only three out of eight bat cell lines were transduced by the H17-VSV pseudotypes, originating from two *Miniopterus* and one *Pteropus* species. This suggests that these viruses may be restricted to a particular set of closely related species belonging to the *Miniopteridae* and *Pteropidae* families, which are closely related to the *Phyllostomidae* family, from which the original H17 and H18 samples were isolated (Agnarsson et al. 2011; Tong et al. 2012). However, this raises further questions, as the cell lines derived from *Pteropus dasymallus yayeyamae* and *Rousettus leschenaultiii* were not permissive, indicating a complex pattern of susceptibility of species to these new influenza viruses. Nevertheless, as these cell lines have only recently been isolated and immortalised (Maeda et al. 2008; Maruyama et al. 2014), detailed characterisation of their surface proteins and expressed proteases are not yet available, requiring additional research before conclusions can be drawn. Further investigation is required to determine the mechanisms involved for the transduction of canine cells but not HEK293T/17 cells, particularly relating to the expression of putative receptors on permissive cell lines.

**Table 1.** Cell lines used in the production of Bat influenza VSV pseudotypes.

| Cell Line | Species | Tissue: |
| --- | --- | --- |
| *Vero E6* | *Chlorocebus sp.* | Kidney |
| *HEK293* | *Homo sapiens* | Kidney |
| **MDCK** | ***Canis lupus familiaris*** | **Kidney** |
| *SK-L* | *Sus scrofa* | Kidney |
| *QT6* | *Coturnix japonica* | Muscle |
| **Cell Line:** | **Bat species:** | **Tissue:** |
| *BKT1* | *Rhinolophus ferrumequinum* | Kidney |
| *FBKT1* | *Pteropus dasymallus yayeyamae* | Kidney |
| ***YubFKT1*** | ***Miniopterus fuliginosus*** | **Kidney** |
| ***IndFSPT1*** | ***Pteropus giganteus*** | **Spleen** |
| *DemKT1* | *Rousettus leschenaultii* | Kidney |
| *ZFBK11-97* | *Epomophorus gambianus* | Kidney |
| ***SuBK12-08*** | ***Miniopterus schreibersii*** | **Kidney** |
| *ZFBS13-75A* | *Eidolon helvum* | Spleen |

List of cell lines used in VSV based pseudotyping of H17 and H18 viruses. Bold entries are those cell lines which were found to be permissive for viral transduction by Maruyama et al. 2016.

**Table 2:** MDCK cell lines.

|  |
| --- |
| NBL-2 (ATCC® cat # CCL-34™) |
| MDCK I (EEACC cat # 00062106) |
| MDCK II (EEACC cat# 00062107) |
| MDCK.1 (ATCC® cat# CRL-2935™) |
| MDCK.2 (ATCC® cat# CRL- CRL-2936™) |
| super dome (ATCC® cat# CRL-2286™) |
| super tube (ATCC® cat# CRL-2285™) |
| Different MDCK cell lines available commercially |

Our results, contrary to previous studies, indicate that N10 is performing a similar function to other NAs in enabling release of new influenza virus particles, which indicate its ability to form VLPs rather than releasing H17 from its cellular receptor (Yondola et al. 2011). PVs bearing H5/H7 and N10 envelope glycoproteins successfully budded from producer cells into the surrounding medium in the absence of a sialic acid cleaving neuraminidase, resulting in significantly increased titres when compared to the same glycoproteins generated without the addition of N10 (Figure 7). This increase in budding PV may be due to the action of N10 on an unknown substrate, or perhaps action of the protein itself in virus morphogenesis and budding (Barman et al. 2004; Yondola et al. 2011). Sialidase activity was however not detected using the enzyme-linked lectin assay, suggesting either a lack of sensitivity of this assay or another mechanism for the removal of surface sialic acids by N10 (Juozapaitis et al. 2014; Sun et al. 2013; Wu et al. 2014). Similarly, PVs bearing solely the N10 glycoprotein did not show sialidase activity via ELLA, or successfully transduce cells to give a significant luciferase reading. Further investigation is required on the role of the bat influenza neuraminidase in its lifecycle, the combination of the findings described in this article, with the fact that the N10 enzymatic structure remains conserved and NA-like, suggests that it is involved in accessory functions other than simply aiding in viral egress. Our results highlight the distinct difference between bat and traditional influenza A viruses where a delicate balance is in place between HA and NA. It may be the case that such a balance exists between bat HA and NAs which will be made clear once the substrate(s) of the bat NA is discovered.

In a previous study, it was demonstrated that the TMPRSS2 protease was capable of inducing HA maturation of H17 through cleavage from HA0 to HA1 and HA2 (Hoffmann et al. 2016). In our study, we demonstrated that this maturation can also be facilitated by TMPRSS4 and HAT. This is of particular interest as the expression of specific proteases is a known limiting factor in viral tropism for a number of different viruses (Böttcher-Friebertshäuser et al. 2010; Ferrara et al. 2012; Millet and Whittaker 2015). This, coupled with the observed susceptibility of a canine-derived cell line to the H17 PV, will need to be factored into future analysis of the potential for zoonotic spillover from bat origin influenza viruses.

The fact that bnmAbs were able to neutralize the virus particles via the H17 hemagglutinin, and their requirement for proteolytic activation, reinforces that we have only scratched the surface in terms of their characterization. Further exploration is required to establish whether these viruses are endogenous or capable of forming infectious particles *in vivo.* Research has revealed that all eight segments of the genome encode functional proteins (Juozapaitis et al. 2014; Moreira et al. 2016; Wu et al. 2014; Zhu et al. 2012), despite the inability to isolate wildtype virus to date. Nevertheless, the development of tools, such as those described here, which can be used in H17 receptor identification studies may ultimately aid discovery of wildtype virus samples from the bat reservoir.

## Methods

### Plasmids

The H17 HA and N10 NA genes from A/little yellow shouldered bat/Guatamala/060/2010 were synthesised commercially by Genscript (Genscript, USA) and subcloned into vector pI.18 (Cox et al. 2002). Lentiviral packaging plasmids p8.91 (Zufferey et al. 1997) and pCSFLW (Demaison et al. 2002) containing a firefly luciferase reporter were used to produce PV. Protease encoding plasmids phCMV-Tag3 (TMPRSS4-myc) and pcDNA3.1-hTMPRSS3 were kindly provided by Prof. Stefan Pöhlmann, Infection Biology Unit, German Primate Center, Germany. pCAGGS-TMPRSS2 and pCAGGS-HAT were kindly provided by Eva Böttcher-Friebertshäuser, Philipps University of Marburg, Germany. The plasmid bearing the Vesicular stomatitis virus envelope protein (VSV-G), pMD.G was obtained from Dr Yasu Takeuchi, University College London, United Kingdom. All the plasmids that were used in this study are listed in Table 3.

**Table 3:**
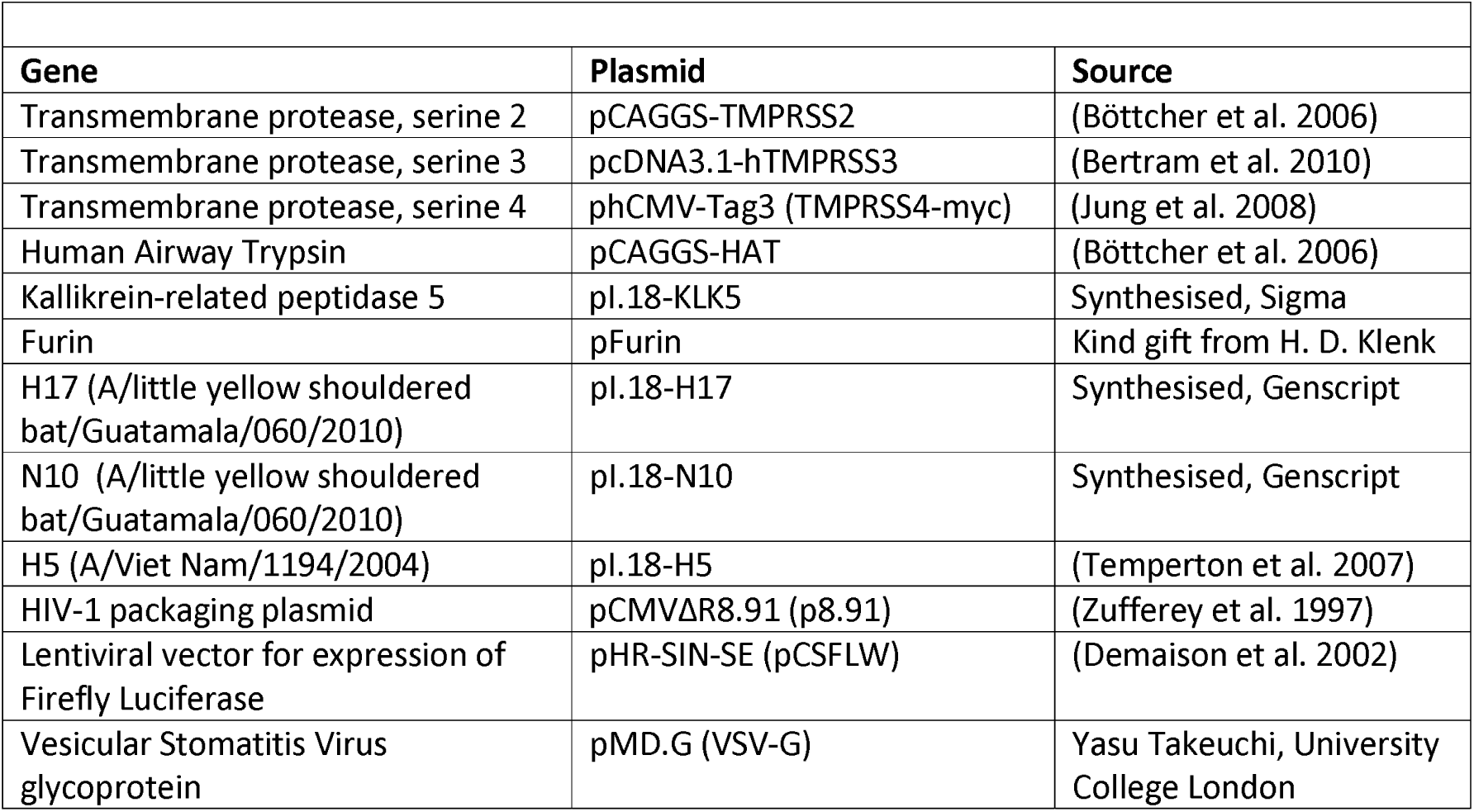
Genes, plasmids and sources.

### Cell lines

Multiple cell lines were used for titration of PV containing supernatants. HEK293T/17 cells were kindly provided by Dr Edward Wright (University of Westminster, UK). MDCK II and RIE1495 were kindly provided by Dr. Gert Zimmer (Institute of Virology and Immunology, Switzerland). U-87 cells were provided by Dr Simon Scott (University of Kent, UK). Madin-Darby Canine Kidney (MDCK) cells were kindly provided by Prof. Sarah Gilbert (Jenner Institute, University of Oxford, UK). All cells were cultured in Dulbecco’s modified eagle medium supplemented with 10% fetal bovine serum.

### Antibodies

mAbs CR9114 and CR6261 were produced by Crucell (Janssen Vaccines AG, Bern, Switzerland). FI6-nt and cholesterol conjugated FI6 (FI6-Chol) were produced by Alfredo Pesci and Krzysztof Lacek from sequence information derived from Corti et al. 2011.

### Production and quantification of H17N10 and H17 bearing lentiviral pseudotypes

PV were produced as described previously (Ferrara et al. 2012; Temperton et al. 2007) and as shown in Figure 1. Briefly, transfection of HEK293T/17 cells was performed using a variety of combinations of plasmids pI.18-H17, pI,18-N10, p8.91 and pCSFLW using polyethylenimine transfection reagent (Sigma Aldrich, UK). Protease-encoding expression plasmids were also included (Table 3). Medium was replaced 12h post-transfection. Supernatants were harvested 48h post-transfection and passed through a 0.45μm filter (Millipore, UK). PV-containing supernatants were titrated using the firefly luciferase Bright-Glo™ system (Promega, UK). Serial dilution of 100μl of PV-containing supernatant was performed across a white flat bottom 96-well Nunclon© plate (Thermo Fisher Scientific, UK). Subsequently, approximately 1×10^4^ cells per well were added per well in 50μl of medium, plates were incubated in a humidified incubator at 37°C 5% CO_2_ for 48h, after which 50μl of Bright-Glo™ substrate was added and luciferase reading recorded in relative luminescence units (RLU) after a 5 minute incubation period. Further sets of N10-bearing PV were produced by transfection of HEK293T/17 cells with 500ng p8.91, 750ng pCSFLW and various amounts of pI,18-N10 plasmid. Medium was replenished after 12h; PV containing supernatants were collected 72h later and passed through a 0.45μm filter. Transfections were carried out in 6-well Nunclon© plates (Thermo Fisher Scientific, UK).

GFP-expressing pseudotypes were produced by substituting the pCSFLW firefly luciferase lentiviral vector mentioned previously with the GFP expressing vector pCSGW. PV-containing supernatants were titrated down clear 96-well Nunclon© plates (Thermo Fisher Scientific, UK) with the addition of 1×10^4^ cells per well. Plates were incubated for 72h and visualized by Nikon Eclipse 50i epiflourescence microscope with a charge-coupled QICAM Fast 1394 camera (QImaging) at x200 magnification.

### Screening of cell lines

Two-fold serial dilutions of PV-containing supernatant were performed as previously described using white 96-well Nunclon© plates (Thermo Fisher Scientific, UK). Subsequently, approximately 1×10^4^ of each cell line was added in 50μl of medium per well. Plates were incubated in a humidified incubator at 37°C and 5% CO_2_ for 48h, after which 50μl of Bright-Glo™ substrate was added and luciferase reading recorded in relative luminescence units (RLU), following a 5 minute incubation period. Control wells were used to measure cell populations for each cell line.

### Pseudotype based microneutralization assay (pMN) using H17N10 and H17 bearing lentiviral pseudotypes

Monoclonal antibodies (mAbs) were serially diluted 1:2 across white 96-well Nunclon© plates (Thermo Fisher Scientific, UK) in 50 μl of DMEM. PV-containing supernatants were diluted and added to each well to give an approximate RLU value of 1×10^6^ per well in 50 μl of DMEM. After 1h incubation at 37°C and 5% CO_2_, approximately 1×10^4^ cells were added per well in a volume of 50μl. Plates were incubated for 48h at 37°C and 5% CO_2_, then 50 μl of Bright-Glo™ was added and luminescence read after a 5 minute incubation at room temperature. Results were analyzed with Graphpad (Prism 7), using nonlinear regression on luminescence values normalized to cell only and virus only control thresholds (100% and 0% neutralization equivalent, respectively). IC_50_ values represent the concentration (ng/ml) required for each antibody to neutralize 50% of functional pseudotyped virus, based on luciferase activity.

### Cell treatment

Overnight treatment of RIE1495 cells with the endosomal acidification inhibitor ammonium chloride (SIGMA) and pretreatment for 2h with neuraminidase (Roche) was done as previously described (Maruyama et.al, 2016). Treated cells were washed three times with serum free medium, and then incubated for 24h with H17 PV. PV titration and luciferase activity was monitored with a luminometer as described previously.

### Pseudotype based enzyme linked lectin assay (pELLA) utilizing N10

ELLA was performed as described by Couzens et al. 2014, but adapted in order to allow use of lentiviral pseudotypes as a source of NA (Biuso et al. 2018; Prevato et al. 2015). Briefly, PV containing supernatant was serially diluted (1:2) across a standard clear microtitre plate in sample diluent (PBS, 1% BSA, 0.5% Tween 20). 50 μl of the resulting dilution series was transferred in duplicate to Nunclon© Maxisorp 96-well plates (Thermo Fisher Scientific, UK) previously coated with Fetuin (Sigma Aldrich, UK) and containing 50 μl of sample diluent per well. Plates were then incubated for 18h and washed, prior to addition of conjugate diluent (PBS, 1% BSA) containing a 1:500 concentration of peanut lectin conjugated to horseradish peroxidase (PNA-HRPO, Sigma Aldrich, UK). Plates were then incubated for 2h in the dark, whereupon OPD-based substrate in citrate buffer was added (Sigma Aldrich, UK). Reactions were stopped after 10 minutes using 50 μl 1M H_2_SO_4_, and readings recorded using a standard ELISA plate reader at 492nm. Exogenous neuraminidase from *Clostridium perfringens* (Sigma Aldrich, UK) was used as a positive benchmark control and samples were assayed alongside PV bearing neuraminidases from other influenza subtypes.

### Statistical analysis

Where possible, statistical analysis was carried out to determine whether differences in PV titre were significant. One-way ANOVA t-tests were performed using fold change scores with a Tukey’s multiple comparisons test. P-values were set at 0.05 (*P* 0.05) unless indicated otherwise. Significantly different data are denoted with asteriks representing p= <0.01 (*), <0.001 (**), <0.0001 (***) and <0.00001 (****).

## Acknowledgements

EG is supported by the Biotechnology and Biological Sciences Research Council (http://www.bbsrc.ac.uk) via Strategic LoLa grant BB/K002465/1 “Developing Rapid Responses to Emerging Virus Infections of Poultry (DRREVIP)”.

